# Acidophilic nitrification alleviates phosphorus deficiency in hydroponics using biogas digestates

**DOI:** 10.1101/2022.12.30.522315

**Authors:** Satoru Sakuma, Ryosuke Endo, Toshio Shibuya

**Affiliations:** Osaka Prefecture University; Osaka Metropolitan University

## Abstract

Biogas digestates can be applied to hydroponics via nitrification. However, the low solubility of phosphorus in digestates can cause phosphorus deficiency in plants. Here, we hypothesized that acidophilic nitrification might prevent this deficiency by dissolving phosphorus in the digestate. Acidophilic and neutrophilic nitrification were conducted at a pH of 3.27 and 6.25 using biogas digestates from food wastes. Acidophilic nitrification dissolved about 3.5 times more phosphorus than neutrophilic nitrification, but the increased acidity also reduced the nitrification rate, resulting in residual ammonium. We then grew lettuce hydroponically with filtrates of these digestates. The growth performance suggested that the increased phosphorus improved growth and that the residual ammonium did not inhibit it. Acidophilic nitrification was shown to be effective for use in hydroponics, particularly to alleviate phosphorus deficiency. These findings should provide new insights into resource recycling, which is essential in both urban and space environments.

**Graphical abstract:** 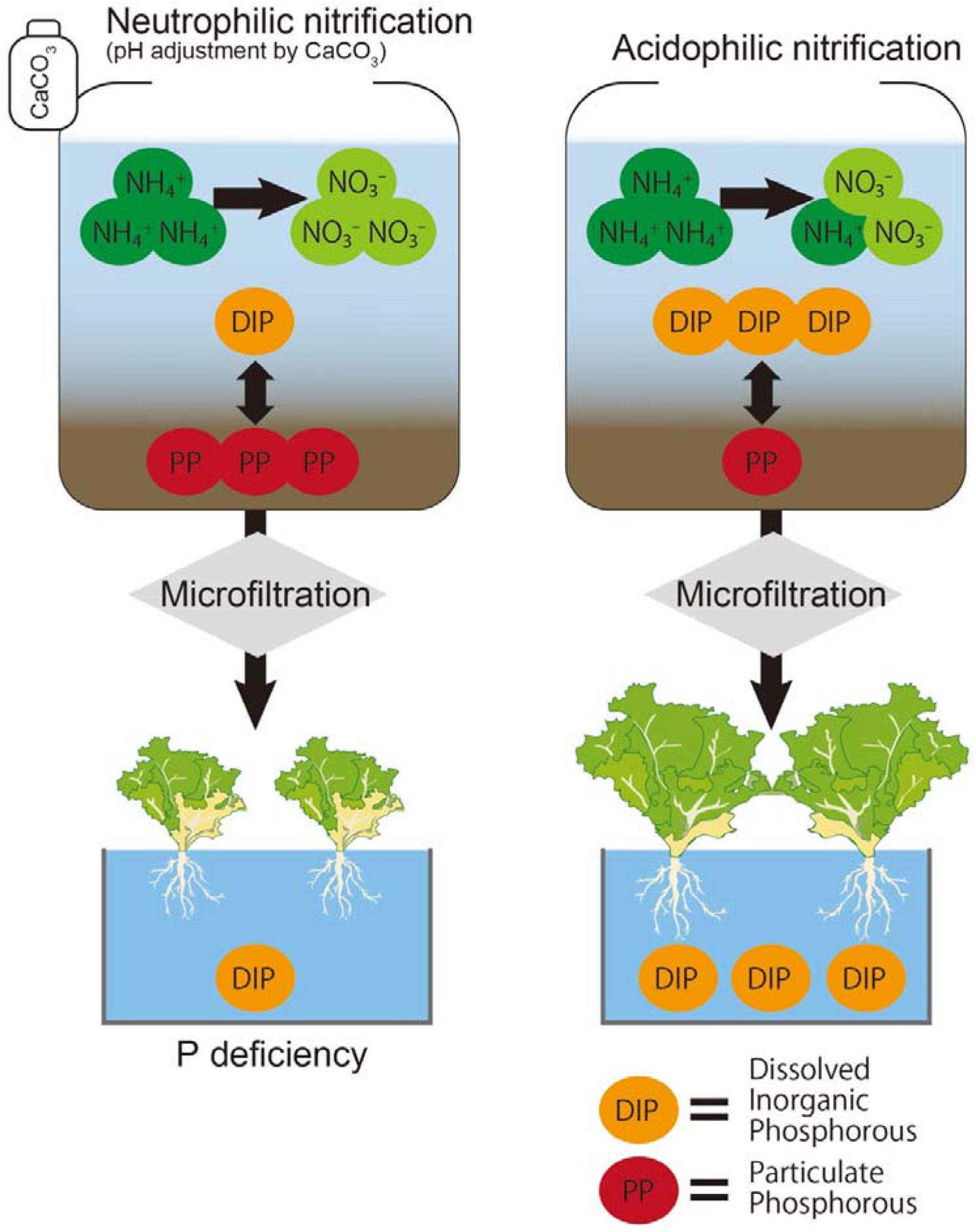

## 1. Introduction

Biogas digestates, residues of anaerobic digestion, can be used in a nutrient solution in hydroponics. Nitrification of the ammonium in biogas digestates enhances this process (Takemura et al., 2020, Lind et al., 2021, van Rooyen and Nicol, 2022) because it helps to avoid the growth inhibition caused by high concentrations of ammonium (Krishnasamy et al., 2012). Our research group has shown that the filtrate of neutrophilic nitrified biogas digestate (f-NNBD) can be used to cultivate chrysanthemum on rock wool (Takemura et al., 2020), but biogas digestates or filtrates of aerobic digestates applied to hydroponics cause phosphorous deficiency (Stoknes et al., 2016, Delaide et al., 2021).

Phosphorus deficiency appears to result from the accumulation of phosphorus in the solid phase of digestates. Three main types of phosphorus exist in activated sludge: inorganic phosphorus, organic phosphorus, and polyphosphate, which account for 10–30%, 5–30%, and 30–80% of total phosphorus, respectively (Yu et al., 2021). Recent studies have shown that acid treatment of activated sludge after biological phosphorus removal enhances the release of phosphorus from the sludge (Latif et al., 2015, Ding et al., 2022) and that the released phosphorus is derived from the degradation of short-chain polyphosphate (Zhang et al., 2021) or the dissolution of inorganic phosphorus (Xu et al., 2015). Therefore, acid treatment is a potentially useful method to avoid phosphorus deficiency because it releases a large fraction of phosphorus in the solid phase.

Acidophilic nitrification, which nitrifies ammonium under acidic conditions, may enhance phosphorus release from the solid phase. Although there is a consensus that the optimum pH for nitrification is between 7 and 8 (Van Hulle et al., 2010), some researchers created sequencing batch reactors that nitrify ammonium from wastewater at a pH of 3–4 (Gisseke et al., 2006, Hanada et al., 2014). Acidophilic nitrification could therefore release phosphorus and alleviate phosphorus deficiency in hydroponics. A potential drawback of this increased acidity, however, is that some ammonium may remain unnitrified because acidic conditions slow the rate of nitrification (Gisseke et al., 2006, Hanada et al., 2014). An increase in ammonium concentration may inhibit plant growth (Britto and Kronzucker, 2002), while the release of phosphorus may simultaneously alleviate phosphorus deficiency.

In this study, to determine whether acidophilic nitrification enhances phosphorus release, biogas digestates were fed into acidophilic and neutrophilic (typical) nitrification reactors, and the digestate compositions were compared. Lettuce was then hydroponically grown using the filtrates of these digestates, and the growth performances were compared between treatments to examine the effect of acidophilic nitrification.

## 2. Materials and methods

### 2.1. Neutrophilic and acidophilic nitrification systems

This experiment used standard food waste as a substrate to perform anaerobic digestion. The biogas digestates were fed into a neutrophilic nitrification reactor and an acidophilic nitrification reactor. The nitrified digestates from the two nitrification reactors were filtered to produce nutrient solutions for hydroponics.

The conventional method of producing a nutrient solution, specifically filtrates of the NNBD (f-NNBD), consists of three processes, as shown in Figure 1a and described by Takemura et al. (2017). In the first process, standard food waste underwent anaerobic digestion in a mesophilic condition at a temperature of 38°C. In the second process, the biogas digestates from the first process were diluted three times with water and fed into a neutrophilic nitrification reactor, which was operated at 28°C and had a hydraulic retention time of 23 days. The pH in the reactor was controlled to approximately 6–7 by adding calcium carbonate when the pH fell below 6. Finally, NNBD extracted from the neutrophilic nitrification reactor was microfiltered using a microfiltration membrane (Microza Lab Module, Asahi Kasei Co., Tokyo, Japan), and the resulting f-NNBD were used in the hydroponics experiments.

**Fig. 1.**
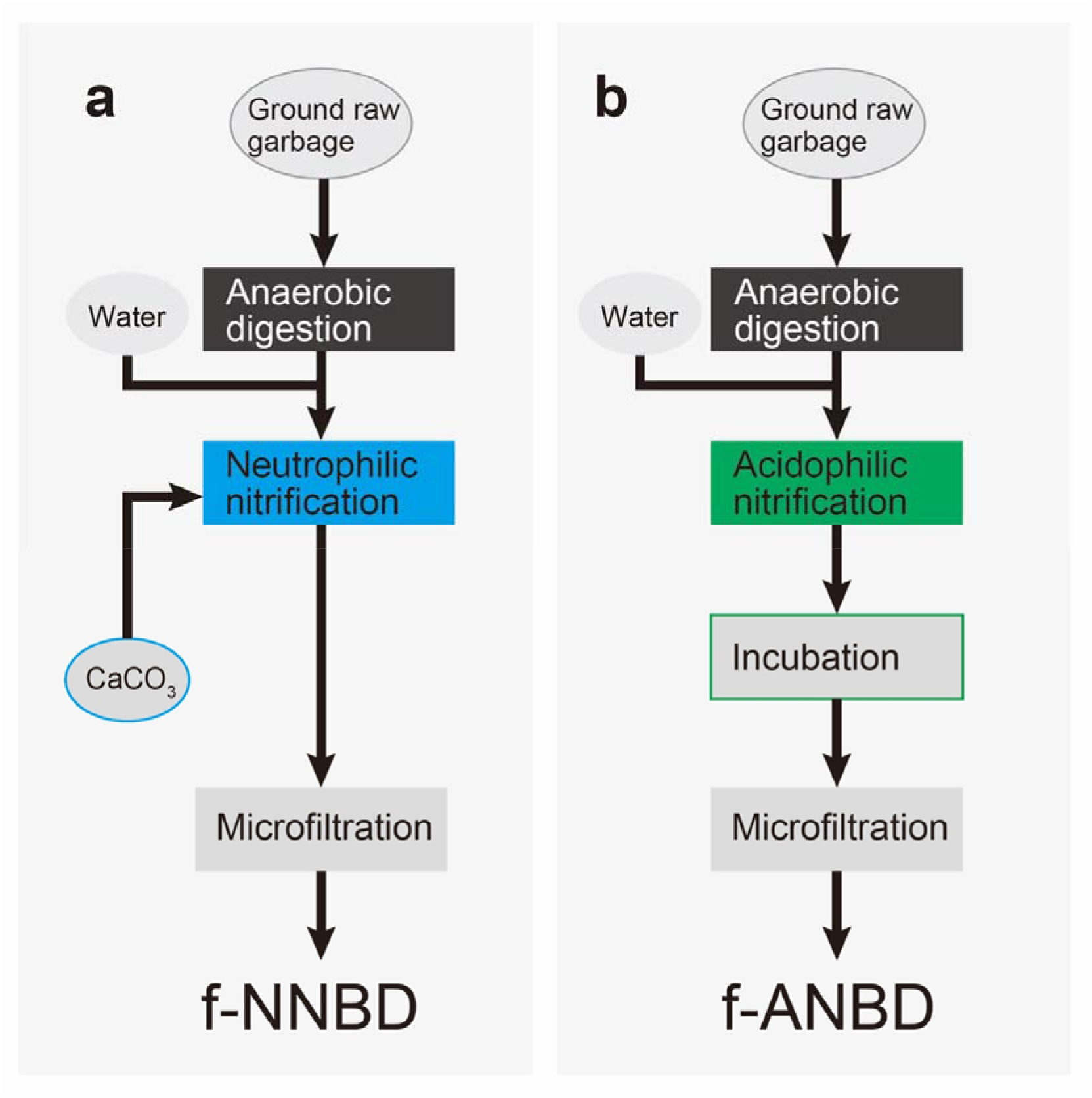
The nutrient solution production system from biogas digestates used in the experiment. Before microfiltration, acidophilic nitrification goes through incubation (see section 2.2). NNBD, neutrophilic nitrified biogas digestate; ANBD acidophilic nitrified biogas digestate; f-NNBD, filtrates of NNBD; f-ANBD, filtrates of ANBD.

The production of acidophilic nitrified biogas digestates (f-ANBD) consists of four processes, as shown in Figure 1b. The first process was identical to the first process of the conventional method, but a different reactor was used. In the second process, an acidophilic nitrification reactor was constructed from the digestates of neutrophilic nitrification. The biogas digestates from the first process were diluted three times with water and fed into the acidophilic nitrification reactor, which was operated at 28°C and had a hydraulic retention time of 45 days. Acidophilic nitrification was performed in an acidic environment by withholding the addition of calcium carbonate, causing the pH to decrease through nitrification. In the third process, approximately 1 L of acidophilic nitrified biogas digestate (ANBD) was withdrawn on a weekly basis, and then it was incubated at room temperature for at least 1 week to increase the pH by denitrification. The pH of ANBD needs to be allowed to increase to about 6 because ANBD is too acidic for use in a nutrient solution. Finally, incubated ANBD was microfiltered in the same manner as NNBD to obtain f-ANBD for use in the hydroponics experiments.

Four independent sets of measurements were performed to analyze the composition of the NNBD, ANBD, and incubated ANBD. Phosphorus and nitrate concentrations were measured by molybdenum blue colorimetry (TNT845, Hach, Loveland, CO, USA) and the dimethylphenol method (TNT836, Hach), respectively, using a spectrophotometer (DR3900, Hach). Total phosphorus was measured after decomposition at 100°C for 1 h using potassium persulfate from a reagent kit (TNT845, Hach). The digestates were centrifuged at 8000 ×*g* for 10 min, and then the supernatants was filtered through a 0.45 μm polyethersulfone membrane to determine the concentrations of dissolved inorganic phosphorus and nitrate. Ammonium ion concentrations were measured by the coulometry method (Quick Ammonia AT-2000, Central Kagaku, Tokyo, Japan). The pH was measured with a compact pH meter (Laqua twin-pH-33, Horiba, Kyoto, Japan).

### 2.2. Lettuce hydroponics experiment

Leaf lettuce *(Lactuca sativa* L. ‘Okayama Salad’) was grown by the deep flow technique using f-NNBD, f-ANBD, and a commercial nutrient solution (OAT House A formula, OAT Agrio, Tokyo, Japan) to investigate the effect of acidophilic nitrification on the growth of lettuce. The nutrient solutions were diluted to an electrical conductivity (EC) of 1.2 mS cm^−1^. Because f-NNBD and f-ANBD contained almost no iron and manganese, ethylenediaminetetraacetic acid iron(III) sodium salt hydrate and manganese chloride were dissolved to 2.7 and 1.5 mg L^−1^, respectively, to match the concentrations in the commercial nutrient solutions. Hydrochloric acid or sodium hydroxide was added to the nutrient solution to maintain the pH at 6.1 before cultivation. The pH of the nutrient solution during cultivation was controlled in the range of 5.7 to 6.3 by a pH control device consisting of a pH sensor (Analog pH Sensor/Meter Pro Kit V2, DFRobot, Shanghai, China), a peristaltic pump (WPM, Welco, Tokyo, Japan), and a microcontroller (Arduino UNO, Arduino, Milan, Italy).

The lettuce seedlings used in this experiment were germinated in tap water after being sown on a sponge and then grown in a commercial nutrient solution for 15 days after germination. Four seedlings of similar size were transplanted into 10 L of f-NNBD, f-ANBD, or commercial nutrient solution, respectively. Lettuce seedlings were cultivated under white LED lights with a photosynthetic photon flux density of 200 μmol m^−2^ s^−1^ at a photoperiod 14:10 h L:D, 22°C, 70% humidity, and 1400 ppm CO_2_ concentration. After the seedlings were raised, each nutrient solution was aerated for 15 days using an air pump. No water was added to the nutrient solutions although the amount of water was reduced by transpiration during the growing period; at least 6600 mL of nutrient solution remained at harvest for all samples.

Nitrate concentration and EC were measured with a compact nitrate ion meter (Laqua twin-pH-33, Horiba) and a compact EC meter (Laquatwin-pH-33, Horiba), respectively. Dissolved inorganic phosphorus concentration in the nutrient solution was measured by molybdenum blue colorimetry using a spectrophotometer (UVmini-1240, Shimadzu, Kyoto, Japan). The elemental composition of each nutrient solution was determined by ICP atomic emission spectroscopy (ICPE-9000, Shimadzu) before and at the end of cultivation. The harvested lettuce was dried at 80°C for at least 3 days, and then the dry matter weight was determined separately for shoots and roots. The elemental composition of lettuce was determined by the same ICP atomic emission spectroscopy following AOAC 985.01 after the dried lettuce was milled.

### 2.3. Statistical analysis

Differences in the growth status of lettuce were analyzed by using Tukey’s HSD test at the 5% significance level. Analysis was conducted in the R statistical software (R Development Core Team, 2022) using the stats package within R. There were four biological replicates in the lettuce grown in each nutrient solution.

## 3. Results and discussion

### 3.1. Acidophilic nitrification

The ANBD was acidic and contained nitrate, whereas the feedstock BD for ANBD contained ammonium but no detectable levels of nitrate were observed (Table 1). These results indicate that nitrification proceeded under acidic conditions to produce ANBD. In addition, the ANBD reactor operated at low pH for more than four times longer than the hydraulic retention time, indicating that acidophilic nitrification produced most of the nitrate in ANBD.

**Table 1.**
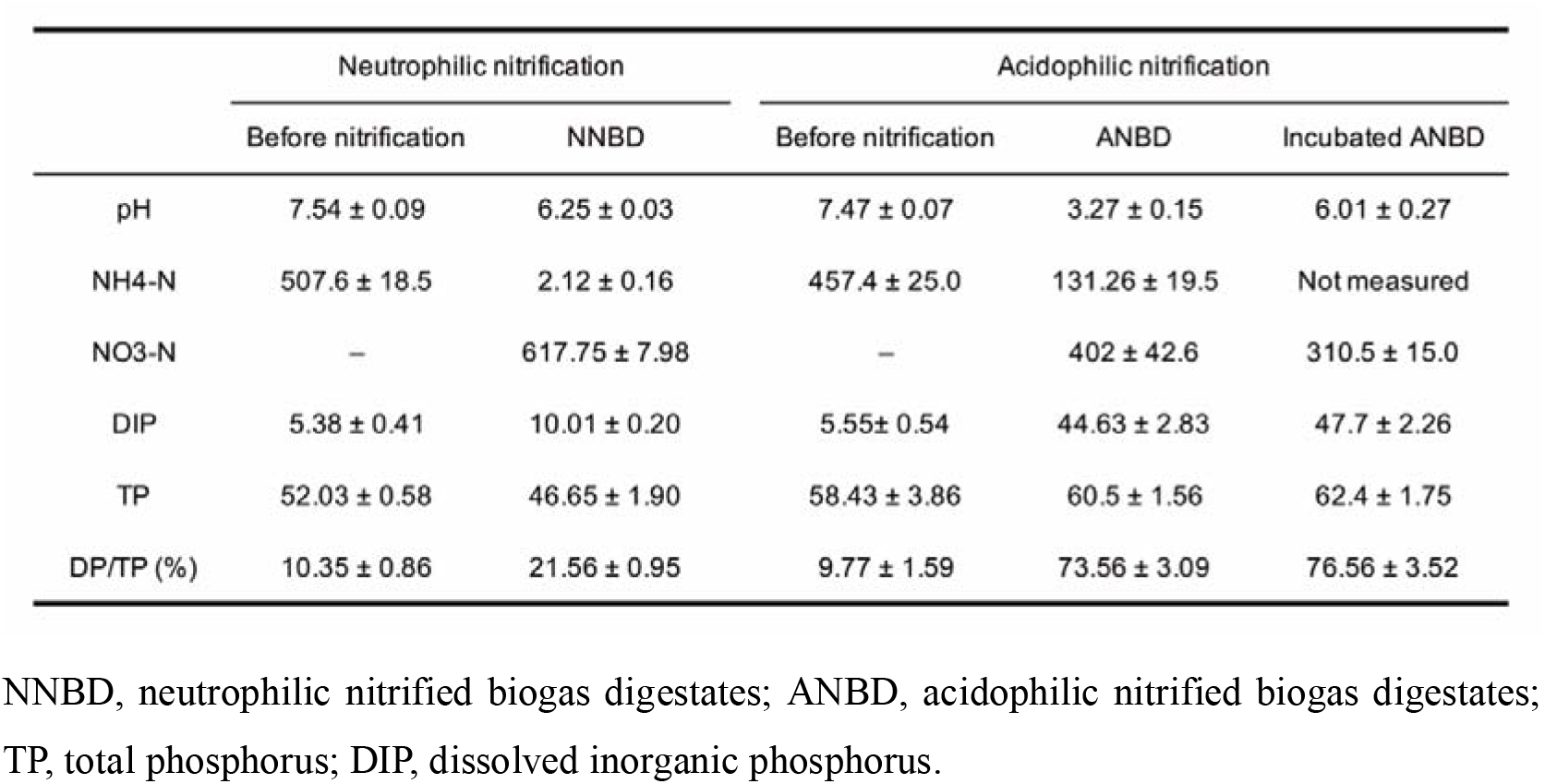
Composition of the digestates in each process. The digestates before nitrification are three times diluted biogas digestates. All units are mg L^-1^ except pH. Values are means ± standard error of four experiments.

The ammonium concentration in ANBD was 60 times greater than that in NNBD (Table 1). The hydraulic retention time of ANBD was about half that of NNBD; therefore, the BD input rate was also about half. In other words, despite the small ammonium load, ANBD did not completely nitrify ammonium. Many studies have shown that acidification inhibits nitrification (Burton and Prosser, 2001, Zhang et al., 2012, Wang et al., 2016). Even for acidophilic nitrification, a pH below 4 decreases nitrification rates (Hanada et al., 2014, Yang et al., 2019). For these reasons, the pH used (3.27) may have caused a higher ammonium concentration in the ANBD via reduced nitrification.

The amount and ratio of dissolved inorganic phosphorus in ANBD were about 4 and 3.5 times larger than those in NNBD (Table 1). In this experiment, the dissolved ratio is more suitable for comparison because the total phosphorus (TP) amount in both digestates differed (Table 1). ANBD was acidic, and NNBD was almost neutral (Table 1). Many studies have reported phosphorus release from activated sludge due to acidification (Latif et al., 2015, Xu et al., 2015, Zhang et al., 2021, Ding et al., 2022). Our results indicate that the low pH of the ANBD notably enhanced the phosphorus release.

The pH of the incubated ANBD increased to an appropriate level for use in hydroponics (Table 1). Moreover, the percentage of dissolved phosphorus in the incubated ANBD did not decrease with increasing pH (Table 1). These results suggest that all the dissolved phosphorus in the ANBD can be available as a nutrient for hydroponics. Notably, neither the low pH of the ANBD nor the pH increase by incubation was due to the application of acidic or alkaline agents, thereby avoiding the cost of chemical application and reducing overall operating costs. The production of a nutrient solution by acidophilic nitrification also has the advantage that no additional chemical inputs are required.

### 3.2. Analysis of lettuce growth

The phosphorus concentration in f-ANBD was one quarter that of a commercial solution and four times that of the f-NNBD (Table 2). This difference between f-ANBD and f-NNBD must be because only dissolved phosphorus passes through microfiltration. The greater amount of phosphorus in f-ANBD potentially complements phosphorus in nutrient solutions. The ammonium concentration in f-ANBD was about twice that of a commercial solution and 80 times that of f-NNBD (Table 2). In contrast to the increased phosphorus concentrations, the increase in ammonium concentrations in f-ANBD could potentially inhibit plant growth.

**Table 2.** Composition of the nutrient solutions used for growing lettuce. All units are in mg L^-1^ except for electrical conductivity (mS cm^-1^).

|  | f-NNBD | f-ANBD | Commercial solution |
| --- | --- | --- | --- |
| EC (mS cm<sup>-1</sup>) | 1.2 | 1.2 | 1.2 |
| pH | 6.1 | 6.1 | 6.1 |
| P | 1.78 | 6.25 | 24.2 |
| NH<sub>4</sub>-N | 0.36 | 25.25 | 12.99 |
| NO<sub>3</sub>-N | 120 | 100 | 98 |
| K | 73.1 | 82.3 | 131 |
| Ca | 168 | 94.8 | 81.2 |
| Mg | 5.61 | 6.5 | 17.4 |

The fresh shoot weight of lettuce grown in f-ANBD was significantly larger than that grown in f-NNBD and equivalent to that of commercial nutrient solution (Fig. 2a), indicating that f-ANBD significantly promoted lettuce growth as compared with f-NNBD. The phosphorus content of shoots on a dry matter basis was significantly greater in the order of lettuce grown in the commercial nutrient solution, f-ANBD, and f-NNBD (Fig. 2b). According to Burns (1987), lettuce growth rate declined when the phosphorus content decreased to 3.3 mg g DW^-1^. Lettuce grown in f-ANBD might have contained sufficient phosphorus to increase dry matter weight, whereas the phosphorus content was too low in f-NNBD and the growth rate was therefore lower. Moreover, the dry matter content and root/shoot ratio of lettuce grown in f-NNBD were both significantly higher than lettuce grown under the other conditions (Fig. 2c, d). Buwalda and Warmenhoven (1999) also reported that the dry matter content and root/plant ratio of lettuce increases with phosphorus deficiency. These observations show that lettuce grown in NNBD potentially suffers from a phosphorus deficiency leading to poor growth, whereas the phosphorus supply by ANBD promotes lettuce growth.

**Fig. 2.**
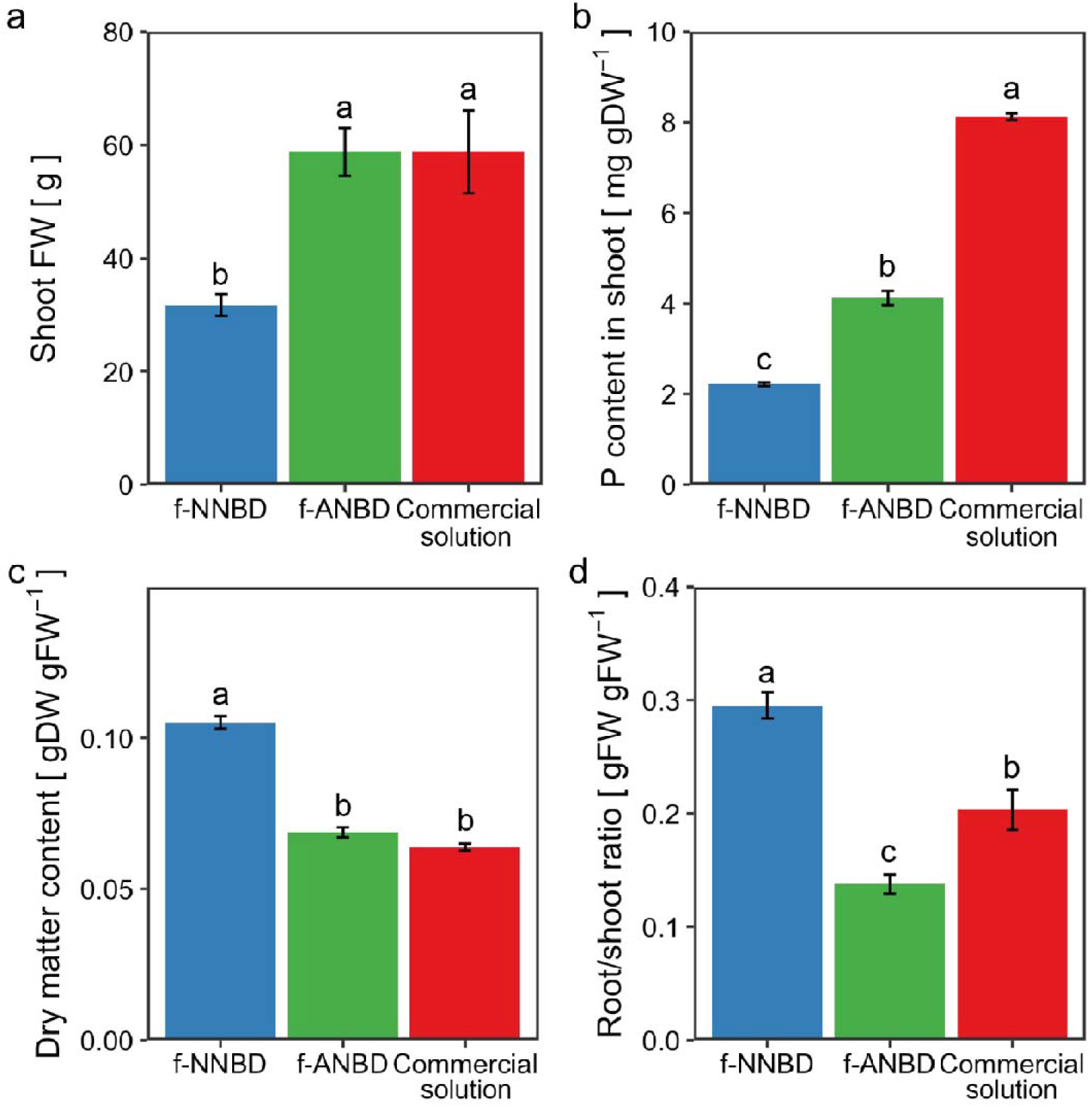
(a) Fresh weight, (b) phosphorus content of shoots, (c) dry matter content, and (d) root/shoot ratio of lettuce hydroponically grown in three different nutrient solutions. Values are mean ± SE (*n* = 4). Different letters within each panel indicate significant differences between the nutrient solutions (Tukey’s HSD test, *p* < 0.05). FW, fresh weight; DW, dry weight; f-NNBD, filtrates of neutrophilic nitrified biogas digestates; f-ANBD, filtrates of acidophilic nitrified biogas digestates.

Chlorosis occurred in lettuce grown in f-NNBD and f-ANBD on day 15 at harvest, but it was not present on day 11 (Fig. 3). Because both nutrient solutions contained almost no phosphorus by day 11 (Fig. S1), phosphorus deficiency could cause chlorosis in both treatments after day 11. Although the symptoms were similar, the area of chlorosis in the lettuce grown in f-ANBD was relatively small (Fig. 3). Therefore, ammonium in f-ANBD did not appear to adversely affect the growth rate or cause chlorosis, which occurs in plants grown in nutrient solutions with high concentrations of ammonium (Britto and Kronzucker, 2002). These results show that the relatively high amount of phosphorus in f-ANBD promoted lettuce growth and alleviated chlorosis, and the greater amount of ammonium caused no observable negative effects on lettuce growth.

**Fig. 3.**
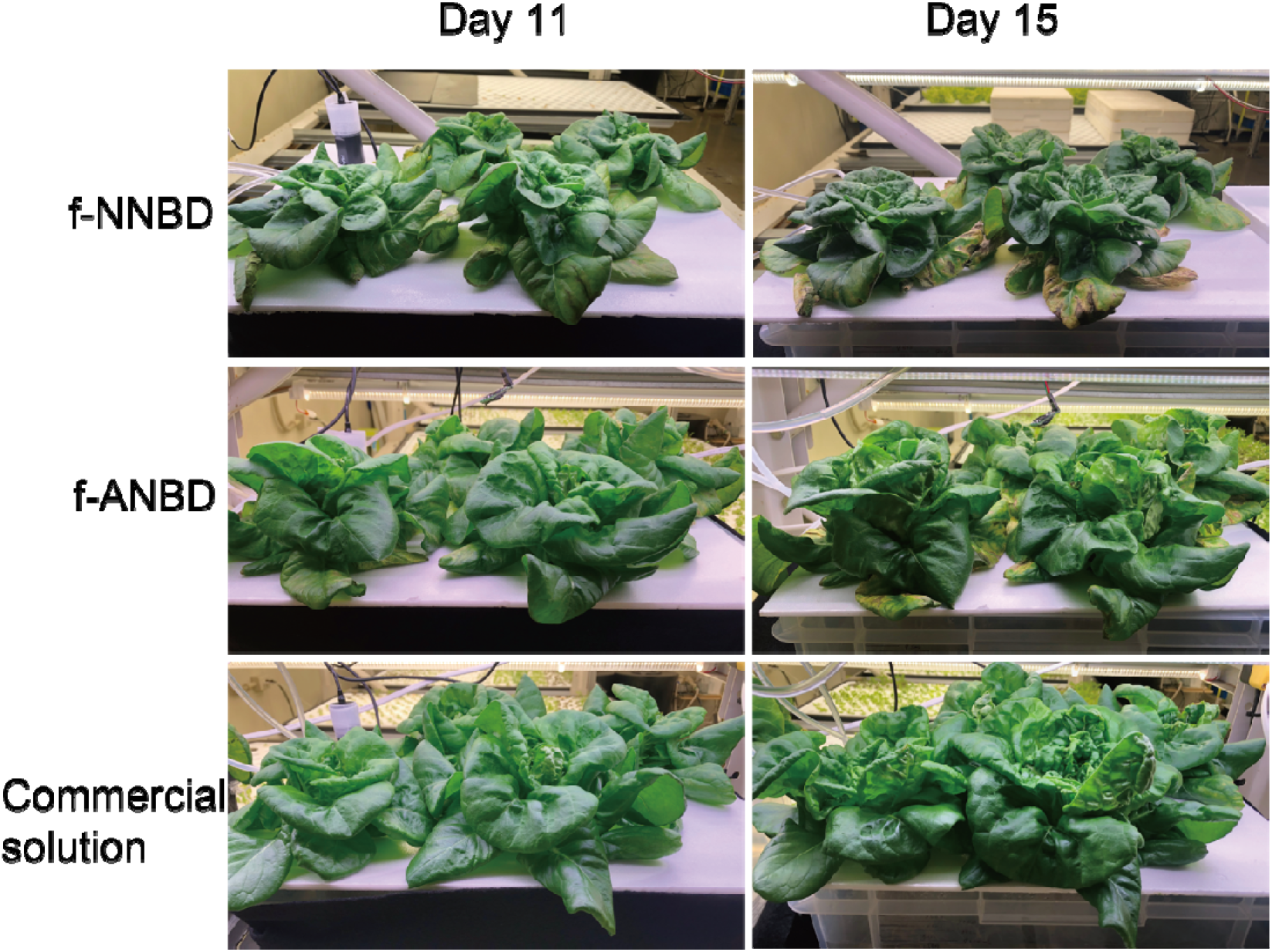
Photographs of lettuce on days 11 and 15 (the day of sampling) following the initiation of the experiment. Lettuce seedlings were cultivated using the three nutrient solutions depicted in Table 2. f-NNBD, filtrates of neutrophilic nitrified biogas digestates; f-ANBD, filtrates of acidophilic nitrified biogas digestates.

## 4. Conclusion

About 3.5 times more phosphorus was dissolved by acidophilic nitrification than neutrophilic nitrification, but the lower pH (increased acidity) slowed the nitrification rate, resulting in more residual ammonium. The fresh weight of lettuce hydroponically grown in the acidophilic nitrification filtrate was twice that of lettuce grown in the neutrophilic filtrate. The growth performance suggested that the increased amount of phosphorus improved growth and that the residual ammonium did not inhibit it. Consequently, we conclude that acidophilic nitrification is effective for use in hydroponics, particularly to alleviate phosphorus deficiency. These findings should provide new insights into resource recycling, which is essential in urban and space environments.

## CRediT authorship contribution statement

**Satoru Sakuma**: Experimentation, Data curation, Conceptualization, Writing – Original draft, Visualization, Validation, Editing, Investigation. **Ryosuke Endo**: Supervision, Writing–Review & Editing. **Toshio Shibuya**: Supervision, Writing– Review & Editing.

## Data availability

Data will be made available on request.

## Acknowledgements

This work was supported by JSPS KAKENHI Grant Number JP20K06350 and JST, the establishment of university fellowships towards the creation of science technology innovation, Grant Number JPMJFS2138. The authors also thank the reviewers for their valuable suggestions and comments on the manuscript.

---

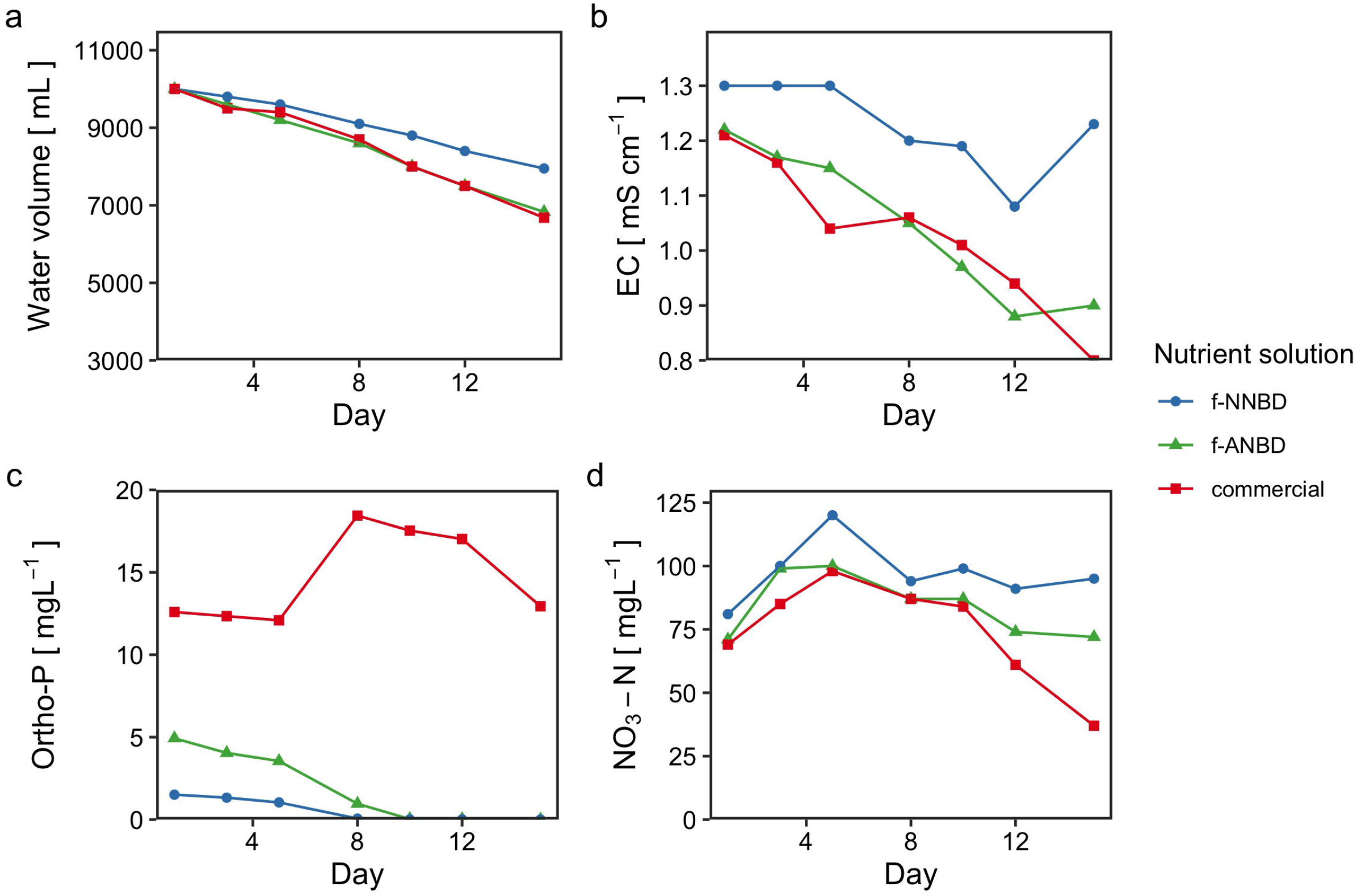

