## Supplemental Table 1 for "Acidophilic nitrification alleviates phosphorus deficiency in hydroponics using biogas digestates"

^a^ *Graduate School of Life and Environmental Science, Osaka Prefecture University, Gakuen-cho 1-1, Naka-ku, Sakai-shi, 599-8531, Osaka, Japan*

^b^ *Graduate School of Agriculture, Osaka Metropolitan University, Gakuen-cho 1-1, Naka-ku, Sakai-shi, 599-8531, Osaka, Japan*

*Sakuma Satoru, Graduate School of Life and Environmental Science, Osaka Prefecture University, Gakuen-cho 1-1, Naka-ku, Sakai-shi, 599-8531, Osaka, Japan*

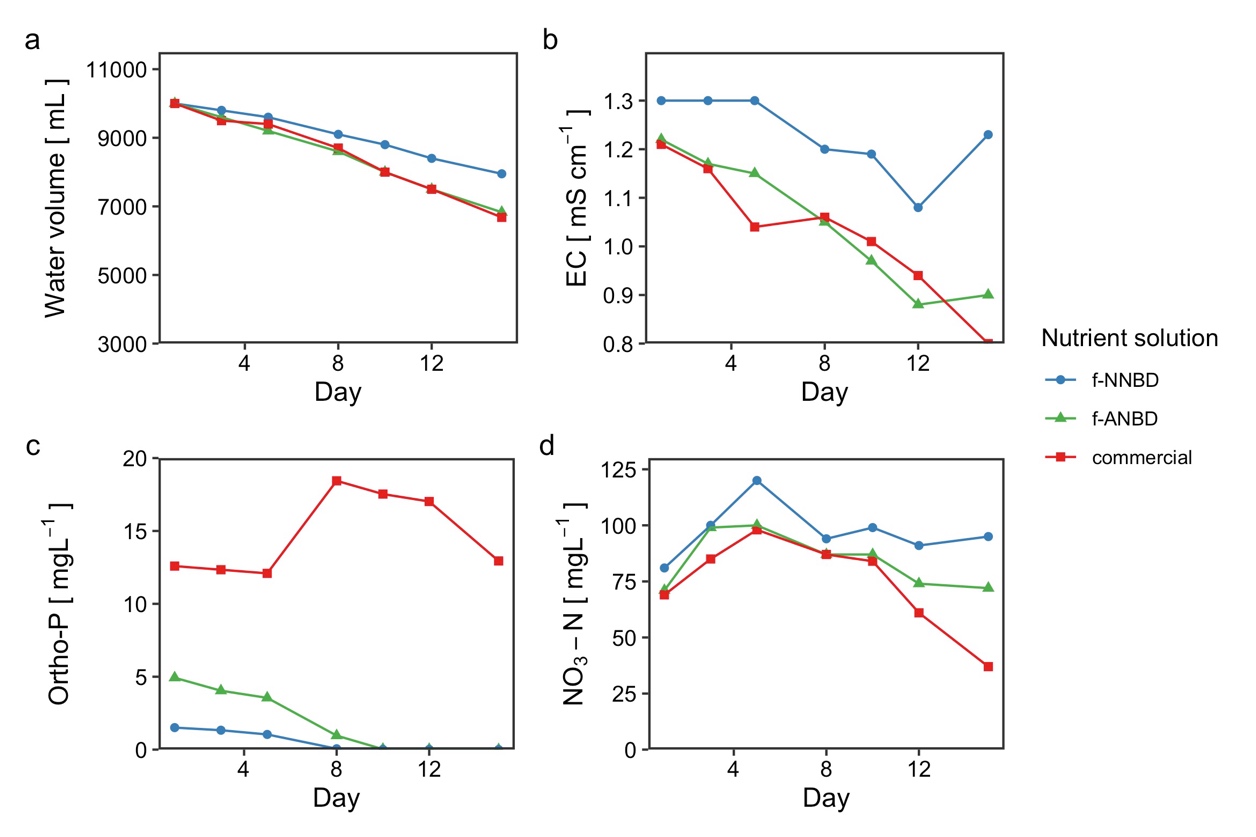

**Fig. S1.** (a) Water volume, (b) electrical conductivity, (c) orthophosphate, and (d) nitrate concentration of each nutrient solution during the experiment. f-NNBD, filtrates of neutrophilic nitrified biogas digestates; f-ANBD, filtrates of acidophilic nitrified biogas digestates.
